# Gaussian embedding-based functional brain connectomic analysis for amnestic mild cognitive impairment patients with cognitive training

**DOI:** 10.1101/779744

**Authors:** Mengjia Xu, Zhijiang Wang, Haifeng Zhang, Dimitrios Pantazis, Huali Wang, Quanzheng Li

**Affiliations:** McGovern Institute for Brain Research, Massachusetts Institute of Technology, Cambridge, MA 02139, USA; Massachusetts General Hospital and Harvard Medical School, Boston, MA 02114, USA; Peking University Institute of Mental Health (Sixth Hospital), Beijing 100191, China; National Clinical Research Center for Mental Disorders & Key Laboratory of Mental Health, Ministry of Health, Peking University, Beijing 100191, China; Beijing Municipal Key Laboratory for Translational Research on Diagnosis and Treatment of Dementia, Beijing 100191, China

## Abstract

Identifying heterogeneous cognitive impairment markers at an early stage is vital for Alzheimer’s disease diagnosis. However, due to complex and uncertain brain connectivity features in the cognitive domains, it remains challenging to quantify functional brain connectomic changes during non-pharmacological interventions for amnestic mild cognitive impairment (aMCI) patients. We present a new *quantitative* functional brain network analysis of fMRI data based on the multi-graph unsupervised Gaussian embedding method (MG2G). This neural network-based model can effectively learn low-dimensional Gaussian distributions from the original high-dimensional sparse functional brain networks, quantify uncertainties in link prediction, and discover the *intrinsic* dimensionality of brain networks. Using the Wasserstein distance to measure probabilistic changes, we discovered that brain regions in the default mode network and somatosensory/somatomotor hand, fronto-parietal task control, memory retrieval, and visual and dorsal attention systems had relatively large variations during non-pharmacological training, which might provide distinct biomarkers for fine-grained monitoring of aMCI cognitive alteration.

---

Azheimer’s disease (AD) is a neurodegenerative brain disorder and the most common form of dementia^1^. However, there is still no cure and no effective drug treatment for AD^2^. Hence, non-pharmacological cognitive intervention for patients at early stages of AD has received a lot of attention due to its non-invasive manner, safety and scalability. Recent studies show that non-pharmacological cognitive intervention can play a positive role in delaying the process or even reducing the cognitive decline for both healthy controls^3^ and amnestic mild cognitive impairment (aMCI) patients^4^. In particular, aMCI is a vital prodromal state of AD harboring memory impairment and has a high risk to develop AD^5^. Multi-domain interventions targeting memory and non-memory domains simultaneously are urgently needed for an optimal aMCI intervention effect. However, most of the previous multi-domain cognitive intervention studies, e.g., the Finnish Geriatric Intervention Study to Prevent Cognitive Impairment and Disability (FINGER)^6^, the French Multidomain Alzheimer Preventive Trial (MAPT)^7^, the Dutch Prevention of Dementia by Intensive Vascular Care (Pre-DIVA)^8^, the Drug and Alcohol Intervention Service for Youth (DAISY)^9^, etc., evaluated the intervention outcomes based solely on neuropsychological assessment or simple characterization of brain anatomical structural changes (e.g., gray matter volume, cerebral ventricle volume). There is a critical need to develop a patient-specific *quantitative* analysis for the underlying *functional* brain regional activity changes during the multi-domain cognitive intervention process. Functional brain network analysis for AD intervention studies can offer great qualitative and quantitative insights into the brain micro-circuits alterations for MCI patients, and could also play an important role in the accurate prediction of the AD progression.

In the present study, we focused on functional brain network analysis for aMCI patients, who completed a multi-domain cognitive training (MDCT) intervention that was designed at the PKU-sixth hospital of China. For each of 12 patients, resting-state functional MRI scans and cognitive assessment scores (MMSE^10^ and MOCA^11^) were collected before and immediately after a 12-week intervention^4^. Aiming to investigate *quantitatively* the underlying functional brain network changes associated with the MDCT intervention, we propose a new approach based on an unsupervised Gaussian embedding-based functional brain network analysis for resting state fMRI data. This method enables mapping of a brain network into multivariate probabilistic Gaussian distributions so as to detect the underlying link changes of functional brain connectomes after the MDCT intervention. Moreover, it provides *uncertainty estimation* for each node in the latent brain network representational space by performing deep learning-based Gaussian embedding for the weighted brain network computed from pre-processed fMRI data using the functional brain template^12^. We compared the new method, called *Graph2Gauss*^*13*^, against other methods, e.g. *node2vec*^*14*^, and cited related literature in the SI. Most of the existing graph embedding methods focused only on a single and binary graph embedding. However, the human brain network is in the form of a weighted graph. Moreover, presently few works consider Gaussian embedding for multigraphs, yet it is prerequisite for quantitative analysis of multi-subject brain networks before and after MDCT intervention. Hence, in our study, we propose a *multi-graph Gaussian embedding (*MG2G) method for the MDCT intervention dataset of aMCI patients.

## Results

### Graph embedding model training and evaluation

An important application of graph embedding is *link prediction* that quantifies how well a model can predict unobserved edges. In order to evaluate the representational performance of the MG2G method, we carried out link prediction experiments on brain networks computed from resting state fMRI data recorded from 12 aMCI patients before and after MDCT intervention. The brain networks were constructed by computing the Pearson’s correlation coefficient between the fMRI time series of 264 brain regions of interest (ROI) that belong to 14 communities (neural systems) according to the Power et al., 2011 brain atlas ^12^. We split the total edges obtained from the network adjacency matrices into three sets: a training set (85%), a validation set (10%) and a test set (5%). The performance in the validation set in terms of AUC (area under the ROC curve) for different values of embedding size *L* is shown in Fig. 1 for a fixed value of *K*=2; here *K* denotes the maximum distance we consider for finding the *k*-hop neighborhoods.

**Fig. 1.**
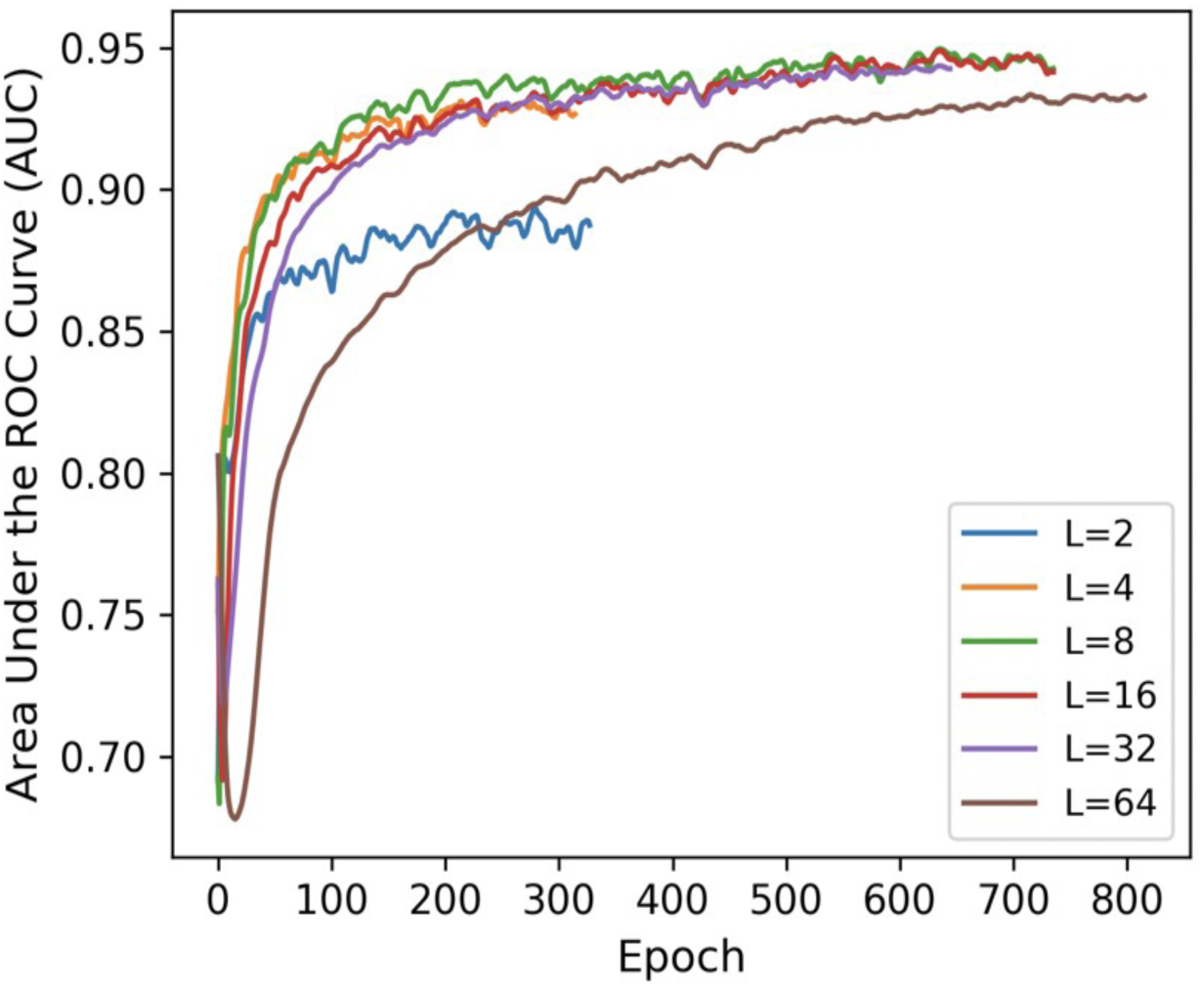
MG2G model performance in link prediction for different values of embedding size (*L*). Results are shown for the validation dataset based on *L* = 2, 4, 8, 16, and 32, with *K*=2 (k-hop neighborhoods).

MG2G achieved high AUC performance in link prediction for embedding size *L* equal to 4, 8, and 16. In contrast, the AUC performance was low for *L* = 2 because embeddings with small size cannot sufficiently capture the representational information of the original graph data. Performance was also low when *L* increased to 32 because an embedding size larger than the latent dimension of the graph may include higher levels of noise. High values of L are also not desirable because they increase computational cost. In addition to evaluating the sensitivity to different embedding size values (*L*) in link prediction, we also evaluated the performance of MG2G for different k-hop neighborhoods (*K*=2 vs. *K*=3); the results are shown in the Supplementary Figure 5 indicating that *K* = 2 is adequate. Finally, for the test set we obtained AUC value of 0.945 in link prediction for a fixed embedding size *L*=2 and k-hop neighborhood *K*=2.

### Quantification of intervention-related brain network alterations using the Wasserstein distance

By performing Graph Gaussian embedding for all patients’ brain networks, every brain region (node) is represented by multivariate Gaussian distributions in a latent space. In order to assess the complex functional network alteration patterns within each patient, we quantified how each node moved in the latent space following the intervention. Specifically, we measured the distances of *each patient’s* brain network embeddings (or Gaussian distributions) for each ROI before and after intervention. The distance measure relied on the 2-Wasserstein distance (W2), which quantifies distances between Gaussian probability distributions. Since our dataset lacks a control group and W2-distance is positive without a known parametric distribution, there is no obvious parametric or non-parametric statistical procedure to apply to these results. However, in the next section we will provide largely consistent results with an alternate group-level analysis.

The within-subject W2 distances for each of the 12 patients are shown in Fig. 2a, with the 264 ROIs and related 14 systems in the brain atlas^12^ described in the Supplementary Table 1. We observe across different patients that the ROI IDs from 112 to 138 exhibited large variations before and after intervention among most patients, and most prominently for subject 3, 6 and 9. Based on the system information from SI Table 1, these regions mainly fall into three functional systems: *default mode, memory retrieval, and visual systems*. Moreover, subject 0 had the greatest number of ROIs with large W2-distance between intervention, and we note that this patient was also diagnosed with depression symptom. There were also patients with smaller variations, namely subjects 4 and 8, compared to other aMCI patients after intervention.

**Table 1.** Names of ROIs with significant network alterations (significant RI index) for each brain system. System name abbreviations same as in Fig. 3.

| System | Significant ROI List (Quantity) |
| --- | --- |
| SSH | Inferior Parietal Lobule(1), Medial Frontal Gyrus(3), Paracentral Lobule(1), Postcentral Gyrus(6), Precentral Gyrus(4), undefined(1) |
| SSM | Precentral Gyrus(2) |
| CoTC | Cingulate Gyrus(1), Insula(2), Medial Frontal Gyrus(1), Middle Frontal Gyrus(1), Superior Temporal Gyrus(1) |
| Audit | Precentral Gyrus(1), Superior Temporal Gyrus(1) |
| DMN | Angular Gyrus(1), Anterior Cingulate(1), Cingulate Gyrus(1), Inferior Frontal Gyrus(1), Medial Frontal Gyrus(3), Middle Temporal Gyrus(6), Parahippocampa Gyrus(1), Posterior Cingulate(2), Precuneus(1), Superior Frontal Gyrus(4) |
| MemRt | Cingulate Gyrus(1), Precuneus(2) |
| Vis | Cuneus(3), Inferior Occipital Gyrus(2), Lingual Gyrus(2), Middle Occipital Gyrus(2), Parahippocampa Gyrus(1), Sub-Gyral(1) |
| FpTC | Inferior Frontal Gyrus(2), Inferior Parietal Lobule(4), Middle Frontal Gyrus(7), Middle Temporal Gyrus(1), Superior Parietal Lobule(1) |
| Sal | Anterior Cingulate(2), Cingulate Gyrus(1), Extra-Nuclear(1), Inferior Frontal Gyrus(1), Middle Frontal Gyrus(3), Sub-Gyral(1), Superior Frontal Gyrus(1), Supramarginal Gyrus(1), undefined(1) |
| SubCt | Extra-Nuclear(3), Thalamus(1) |
| VenAtt | Inferior Frontal Gyrus(2), Inferior Parietal Lobule(1), Superior Frontal Gyrus(1), Superior Temporal Gyrus(2) |
| DorsAtt | Middle Frontal Gyrus(2), Middle Temporal Gyrus(1), Sub-Gyral(1), Superior Parietal Lobule(2) |
| Uncert | Culmen(1), Fusiform Gyrus(1), Inferior Occipital Gyrus(2), Inferior Temporal Gyrus(2), Lingual Gyrus(2), Middle Frontal Gyrus(1), Sub-Gyral(1), Superior Frontal Gyrus(1), Uncus(2) |

**Fig. 2.**
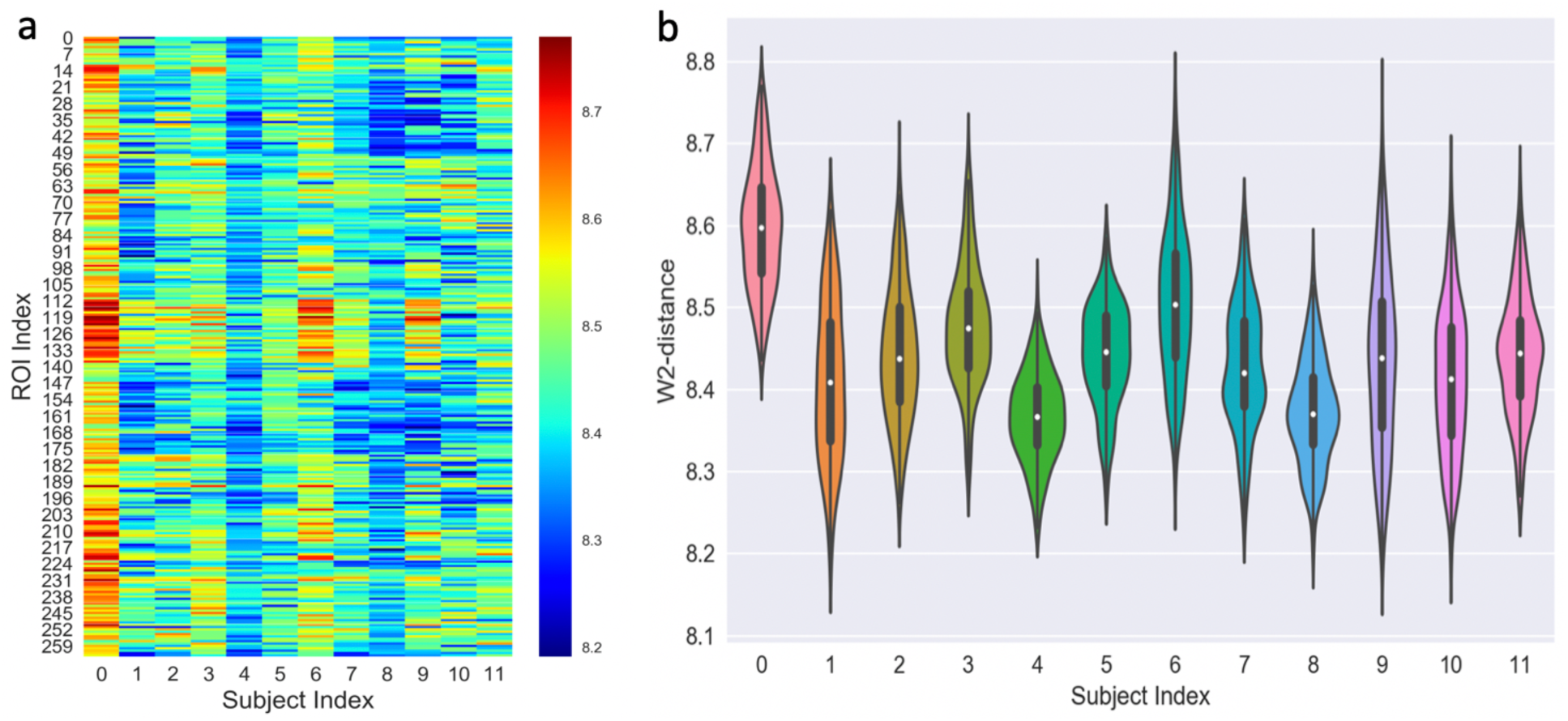
Within-subject intervention-related brain network alterations. (a) W2-distance before and after intervention for each of the 264 ROIs across the 12 patients (*L*=16, *K*=2). (b) Violin plots of the W2-distance distribution over the 264 regions for each of the 12 patients.

To better assess the overall network alteration at the subject-level, we used a “violin plot” (combination of box-plot and density plot) to visualize the W2-distance distributions and probability densities for different patients (Fig. 2b). Each “violin” contains a box-plot (white dot, vertical thick black box and thin black line). The white dot represents the median of W2 distances at each column in Fig. 2a, the vertical thick black box indicates the inter-quartile range, and the thin black line denotes the extensions to the maximum and minimum values. The shaded areas surrounding the box plot show the probability density of the W2-distances across the 264 brain regions for each patient. These results reveal that patients varied considerably with respect to the network alterations, with some subjects exhibiting large W2 medians and variability (e.g., subjects 0 and 6) and others the opposite (e.g., subjects 4 and 8), while there are also some unique subjects with multi-modal shape of the W2 distribution (e.g., subject 7).

To more specifically quantify the ROI-level W2-distance density changes across the 12 patients, we constructed the Kernel Density Estimation (KDE) plot in Fig. 3a. The brain regions (ROIs) around the index ranges of 100-150 and 221-240 exhibited larger alterations (in terms of the W2-distance) compared to other brain regions. As shown in SI Table 1, these regions belong to the following communities (brain systems): *default mode, memory retrieval, visual and dorsal attention*. Additionally, from the KDE plot we can also distinguish three dark blue areas (default mode, visual, frontal-parietal task control, dorsal attention and uncertain) with high probability densities of W2-distance compared to other regions. Subsequently, we obtained the top-15 brain regions for *all patients* measured by the W2-distance, and identified the brain systems they belong to, shown in blue bars in Fig. 3b. Here, the vertical axis denotes the *total* number of top-15 ROIs corresponding to each community. The highest system-level MG2G results identified with this analysis were: *default mode, visual, uncertain, dorsal attention, salience, subcortical, sensory/somatomotor hand*, and *memory retrieval*, which overlap with the dark blue areas in Fig. 3a.

**Fig. 3.**
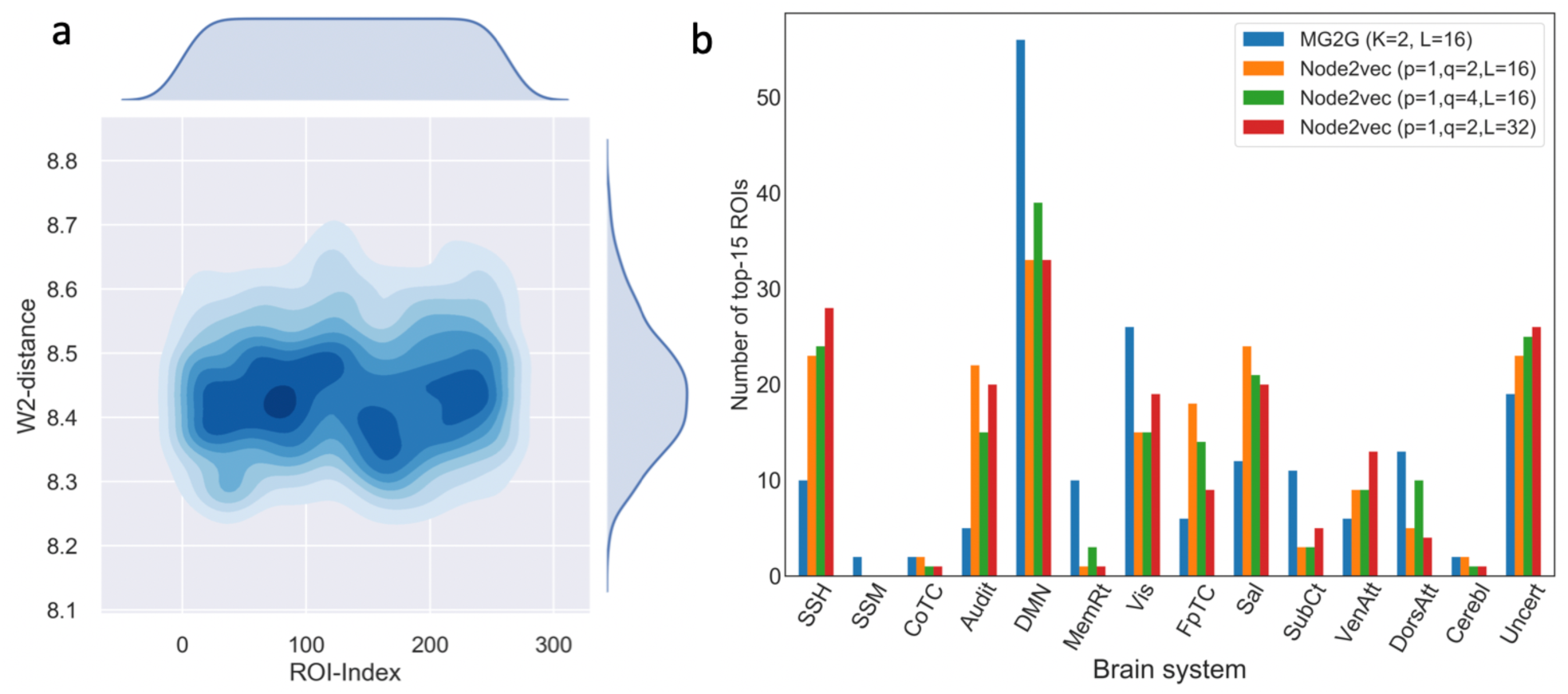
Within-subject intervention-related alterations at ROI-level and system-level. (a) Kernel Density Estimation plot of the W2-distance across all 264 ROIs. (b) Quantification of functional/system-levelb changes for *all* 12 patients before and after MDCT intervention based on MG2G (blue) and node2vec (yellow, green, and red, corresponding to different node2vec parameters). SSH: sensory/somatomotor hand; SSM: sensory/somatomotor mouth; CoTC: cingulo-opercular task control; Audit: auditory; DMN: default mode; MemRt: memory retrieval; Vis: visual; FpTC: fronto-parietal task control; Sal: salience; SubCt: subcortical; VenAtt: ventral attention; DorsAtt: dorsal attention; Cerebl: cerebellar; Uncert: uncertain.

To further validate these system-level results, we also performed a secondary analysis using a different graph-embedding method, the deterministic “node2vec”^14^. The node2vec results are shown in yellow, green and red bars in Fig. 3b; here we performed a similar analysis as in MG2G, but the metric was Euclidean distance because node2vec is deterministic and nodes are mapped to point-vectors in the latent space. We assessed the sensitivity of the results change for different embedding size (*L* = 16, 32) and different hyperparameters (*p* and *q* values) in *node2vec*; these values control the neighborhood exploration in *node2vec*. The *default mode*, sensory/somatomotor hand, auditory, *visual, salience, and uncertain* communities exhibited large subject-level intervention effects. Additional system-level comparisons at the single-subject level can be found in Supplementary Figures 3 and 4 using both the proposed MG2G as well as the node2vec method. We observed some variability among the patients and the two methods (MG2G and node2vec) but overall the top-15 changes in brain regions among the 12 patients mostly occurred in the *default mode, visual, uncertain, salience, memory retrieval, fronto-parietal task control, dorsal attention.*

### Statistical evaluation of intervention-related brain network alterations at the group-level

Here we quantified the intervention-related brain network alterations by defining a new measure, the *reorganization index*, which captured cross-subject W2-distance intervention effects. For every pair of *different* subjects, we computed the W2-distance per ROI when: i) one subject was before and the other after intervention (between-pair), or ii) when both subjects were paired before intervention (within-pair). The former assessed cross-subject intervention-related effects, whereas the latter established a baseline cross-subject W2-distance. We then defined the reorganization index RI as the averaged W2-distance of the between-minus within-pairs. Given 12 patients, we obtained 66 between-pair W2-distances matched by an equal number of within-pair W2 distances, allowing us to perform one-sample t-tests for statistical evaluation.

The between-pair and within-pair distances are exemplified for the ROIs belonging to the ‘Sensory/Somatomotor Hand’ neural system in Supplementary Fig. 6. The between-pair (blue) were largely above the within-pair (red) W2 distances, demonstrating that RI increased due to the intervention for most of the ROIs.

RI results for all 264 ROIs are shown in Fig. 4, with statistically significant results highlighted with red bars (p<0.05, one-sample t-test, false discovery rate corrected). The majority of the ROIs had significantly positive RI, which suggests extensive fMRI brain network reorganization following the MDCT intervention. In Fig. 5, we counted the number of significant ROIs belonging to each neural system. The results indicate that the most extensive brain network reorganization encompassed the *default mode, somatosensory/somatomotor hand, fronto-parietal task control, visual, salience, dorsal attention and uncertain* brain systems. These systems largely overlap with the neural systems identified with the within-subject analysis in the previous section. A list of the significant ROIs contained within each neural system is presented in Table 1.

**Fig. 4.**
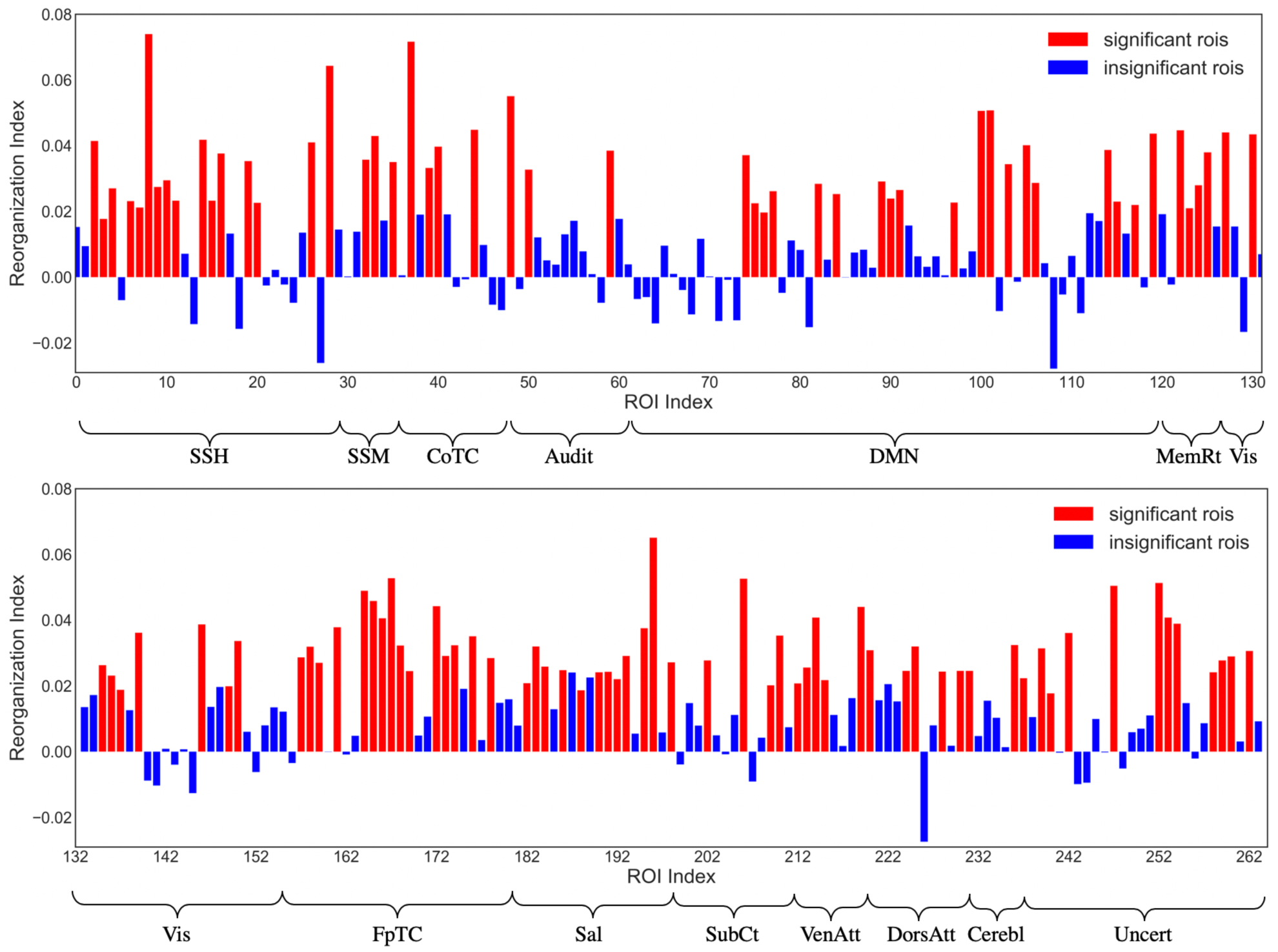
Reorganization index for each of the 264 ROIs. A large number of ROIs had significant RI (red bars; p<0.05, FDR corrected), suggesting extensive intervention-related brain network reorganization. System name abbreviations same as in Fig. 3.

**Fig. 5.**
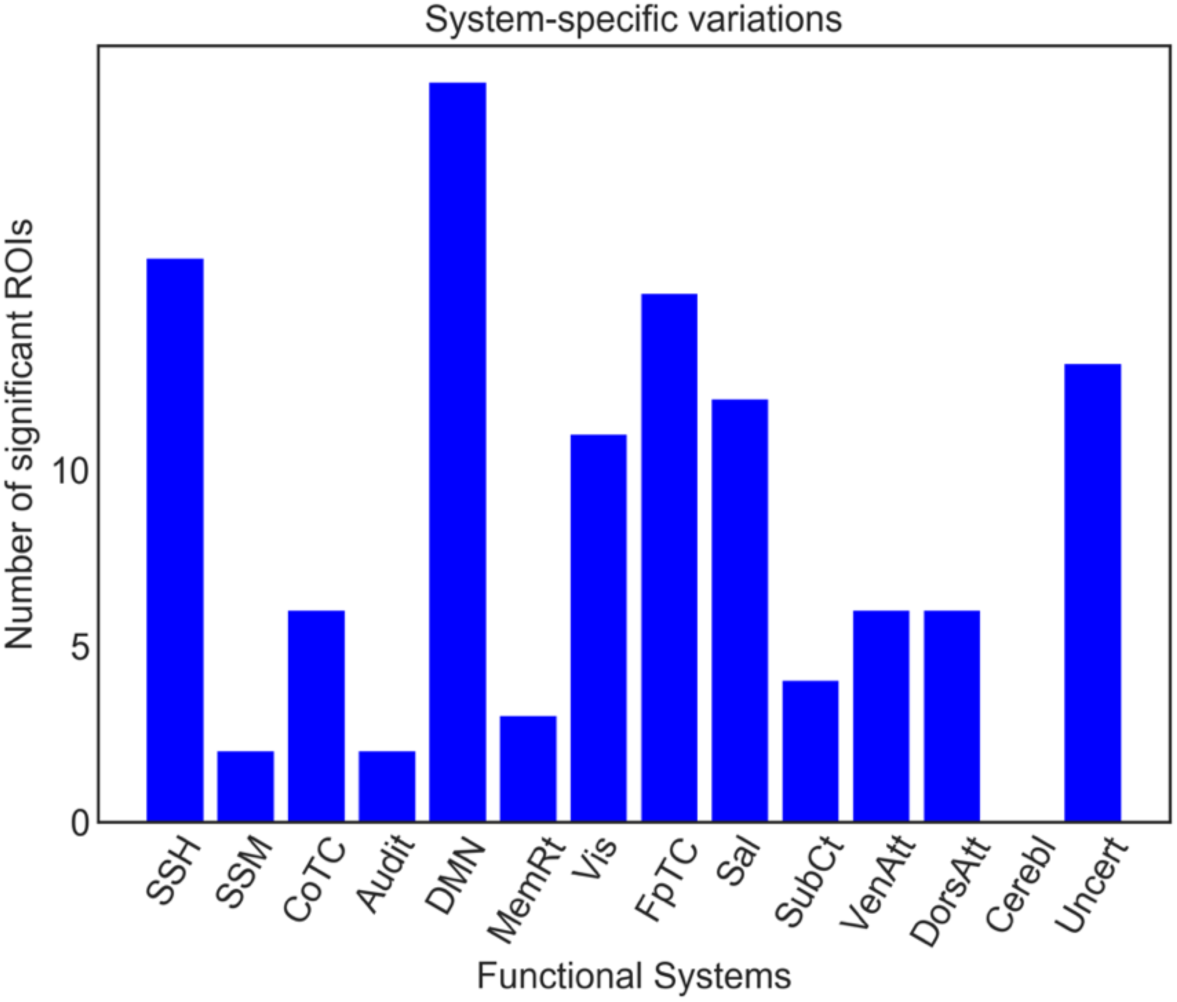
Number of ROIs with significant network alterations (significant RI index) contained within different functional brain systems. System name abbreviations same as in Fig. 3.

### Nodal uncertainty quantification

With Graph2Gauss embedding, every brain region was encoded as a multivariate Gaussian distribution. Hence uncertainty, quantified by the variance, can also be assessed using this graph embedding approach. Fig. 6 illustrates the nodal uncertainty results of graph embedding at baseline and after intervention averaged across all patients. The vertical axis shows the embedding variance for each of the *L*=16 dimensions. Dimensions 8, 10, and 11 had consistently high variance values for the majority of nodes before and after intervention. Dimensions with high uncertainty are unstable and do not contribute to a low-dimensional embedding in the latent space^13^. Thus, we can infer the effective latent dimension to represent our brain network to be equal to (*L –* 3) by excluding the highly unstable dimensions. This yields an effective dimension of 13 (since the embedding dimension was L=16), which is approximately equal with the ground truth community number (14) in the brain atlas. Therefore, our proposed method for fMRI data analysis not only predicted the latent representations, but also yielded the effective dimensionality of the low-dimensional space (latent dimension) by monitoring (during training) the “uncertain” dimensions. More detailed uncertainty quantification results by plotting the corresponding Gaussian distributions are shown in Supplementary Figure 2.

## Discussion

The new method MG2G we introduced, and other recent graph embedding techniques, hold great promise in diverse real-world applications. However, so far the studies incorporating prevalent graph embedding techniques for the analysis of complex and heterogeneous *functional brain network systems* for brain disorders (e.g. Alzheimer’s, Parkinson’s, etc.) are scarce. For example, Rosenthal et al.^15^ first proposed to use a connectome embedding method, node2vec^14^, for the mapping of high-order relations between brain structure and function. As discussed earlier, this method cannot model important uncertainty information about nodal embedding in the latent space. We have applied node2vec in our study to verify the results of MG2G, which in addition can effectively quantify uncertainty for the learned node representations. Therefore, *Gaussian embedding* can facilitate functional brain connectome analytics by employing a stochastic quantitative analysis, which is necessary given the lack of big data and the sensitivity and diversity of the human brain connectomes. To this end, we proposed a new functional brain network analysis framework based on multiple brain connectome Gaussian embeddings via deep neural networks, combined with weighted information of the original graphs. Additionally, we adopted the Wasserstein distance (*W2*) to quantify the brain region (ROI)-level differences between the multivariate embedded Gaussian distributions before and after intervention (Fig. 2a). We constructed violin and KDE plots to estimate and display the W2 distance distributions (Figs. 2b and 3a) from two different perspectives (patient-specific and ROI-specific) and developed a group-level analysis to statistically validate our findings (Figs. 4 and 5). Our results demonstrated that Gaussian embedding-based functional brain network analysis can automatically and quantitatively detect the underlying multiscale (region->system->patient) *subtle changes* of brain networks after non-pharmacological MDCT interventions for aMCI patients. Moreover, we demonstrated two main advantages of the nodal embedding uncertainty in our study: i) we can obtain the intrinsic dimensionality (*L*) of the brain network, and ii) we can quantify the heterogeneity (diversity) of node’s neighbors. The latter is because the high uncertainty to some nodes is due to potential connections with neighbors of different communities with possibly contradicting underlying patterns.

Furthermore, the deep neural network-based model we employed in our study enabled learning the highly non-linear mapping from the original high-dimensional brain network space into low-dimensional Gaussian distributions, while at the same time quantifying the uncertainty about the node embeddings. This is in line with the recent successes of emerging deep learning techniques in diverse fields, when compared to traditional matrix-factorization methods (e.g. SVD^17^) and random walk-based models (e.g., node2vec^14^). Our MG2G model can readily scale up to large-scale network applications unlike traditional methods.

To evaluate the robustness and generalization of the MG2G method, we compared with the *node2vec* method employed in the work of Rosental et al^15^. We compared the two methods in SI (see Supplementary Figures 3 and 4 for details) using the same data as in our main study. Another alternative method is spectral embedding^18^ designed to use an “informative” eigenvector decomposition, however, it becomes inefficient and unstable for large-scale and noisy fMRI data. In contrast, the *node2vec* approach produced comparable results as our proposed MG2G method, as shown in Fig. 3b and in Supplementary Figures 3 and 4, but ignored critical uncertainty information about the node embeddings and the intrinsic system dimensionality. Such information is potentially important for the dynamic, heterogeneous and complex functional role of different regions in the brain connectome. Our proposed deep neural network-based Gaussian embedding model can overcome the aforementioned problems effectively, and obtain *probabilistic* node representations, while preserving both local and global graph topology properties of brain networks.

In addition to the within-subject analysis quantifying network alterations after intervention, we also analyzed statistically network alterations at the group-level by defining a new measure, the reorganization index (RI). In this case too, we found that a large number of ROIs were affected after intervention (Fig. 4), and these changes at the system/community level (Fig. 5) were comparable to the ones we obtained with the within-subject analysis in Fig. 3b. Taken together, our results using two different approaches (MG2G and node2vec) and two different methods of analysis (top-15 ROI and t-test) showed consistency in the regions affected by the MDCT intervention, with details of each region presented in Table 1.

In addition to fMRI networks, in previous work^4^ we have investigated the MDCT intervention effects on structural MRI data and found significant increases in gray matter volume in the right angular gyrus and other subareas following the MDCT intervention. In the current study, we further investigated the underlying MDCT intervention effects at both ROI-level and community-level on the fMRI networks. Therefore, MG2G can provide a more elaborate, cross-modality quantification of network alterations. Specifically, we quantified the differences between probabilistic Gaussian embeddings of functional brain connectomes before and after intervention using the W2-distance metric. The results revealed significant changes on an extensive number of brain regions (Fig 4 and Table 1). Also, system-level changes occurred primarily in the *default mode, somatosensory/somatomotor hand, fronto-parietal task control, memory retrieval, visual and dorsal attention* brain systems (Figs. 3b and 5). Moreover, network alterations varied across patients (Fig. 2), which is consistent with the heterogeneous clinical score profiles.

The broad intervention-related alterations on the intrinsic functional networks may reflect adaptive mechanisms of information integration among different functional systems over the whole brain, due to putative co-activation during the multi-domain training. A previous study that used only explicit-memory training has found increased activation and connectivity in distributed neural networks mediating explicit-memory functions^19^. Hence, an integrated cognitive training that targets more cognitive domains should stimulate more diverse distributed networks underlying multiple cognitive functions. A recent study using MDCT in a healthy older population has found increased functional connectivity within three higher cognitive networks that overlap with our current study: default mode, salience, and central executive network^3^. Therefore, our findings here suggest that widespread changes in functional connectivity induced by MDCT may be due to an enhanced restoration by functional reorganization that benefits brain cognition.

In the future, to better assess and validate the MG2G method on the MDCT intervention study, we plan to extend our method to process multi-modality data (fMRI, MRI, MEG, genetic, and PET) given the multifaced nature of AD. Further, as more subjects enroll in the study and longitudinal data become available, we will better characterize the effectiveness of the MDCT intervention. Specifically, it is important to complete a longitudinal study that facilitates dynamic brain network fluctuation modeling during intervention (*i.e.*, temporal and spatial patterns). Collecting data from a control group will also enable a direct comparison of network alterations across populations for a deeper understanding of the underlying mechanisms of the MDCT intervention.

## Methods

### Participants

All aMCI participants were recruited from the Dementia Care Research Center of Peking University Institute of Mental Health (DCRC-PKUIMH) between May 2015 and September 2015. Twelve of them met the inclusion criteria and completed both a standardized neuropsychological evaluation and MRI scanning at Peking University Third Hospital. All participants were required to be equal to or more than 55 years old, right handed, and have an education level of no less than five years. They were also required to meet the MCI criteria, according to Petersen et al.^20^ as follows: (a) subjective memory complaint, confirmed by an informant; (b) a mini-mental state examination (MMSE) score of no less than 24; (c) an ADL score of no more than 26, and not diagnosed as having dementia (according to ICD-10 and NINCDS-ADRDA criteria). Other inclusion criteria were: a global clinical dementia rating score of 0.5 and no depressive symptoms (Hamilton Depression Scale score ≤12). Exclusion criteria were: a current or past neurological disorder or a current neuro-psychiatric disorder listed in the DSM-IV affecting cognition; currently taking cognitive enhancers; and any physical condition that could preclude regular participation in the intervention program. The present study was approved by the ethics committee of Peking University Institute of Mental Health (Sixth Hospital), Beijing, China. All participants were fully informed regarding the study protocol and provided written informed consent.

### MDCT intervention and cognitive assessment

We used a self-controlled design to investigate the effect of the MDCT program on spontaneous brain activity in older participants with aMCI. Every patient underwent 24 training sessions delivered twice per week over approximately 12 weeks. Each session lasted 60 minutes and included tasks that covered three different cognitive domains. The participants spent 20 minutes engaged in each task per session. The 24-session intervention targeted multiple cognitive domains across the different sessions, including *reasoning, memory, visuo-spatial skill, language, calculation, and attention*. Neuropsychological assessments and MRI scans were conducted before and after the 12-week training program; details are described below.

### Imaging protocol

MRI was performed using a 3T General Electric MRI 750 (Chicago, Illinois, United States) with an 8-channel sensitivity-encoding head coil (SENSE factor = 2.4), with parallel imaging using a Gradient-Recalled Echo-Planar Imaging (GRE-EPI), at the Peking University Third Hospital Neuroimaging Center. Two resting state BOLD fMRI imaging data were collected for each of the 12 aMCI patients, one before and one after MDCT intervention. The resting state functional MRI (rs-fMRI) data in each patient consisted of 230 functional volumes, each slice had a 64 × 64 grid, time repletion (TR) = 2000 ms, time echo (TE) = 20 ms, flip angle= 90°, field of view (FOV)= 240×240 mm^2^, 41 axial slices, thickness = 3.0 mm, spacing between slices = 3.3 mm, acquisition matrix = 64 × 64.

### Cognitive assessment

We applied a comprehensive cognitive test battery to evaluate the cognition of patients at the baseline and after 12-week MDCT intervention. Global cognition was assessed via the MMSE (range 0–30) and MOCA (Montreal Cognitive Assessment) (range 0– 30), with higher MMSE scores indicating higher levels of global cognition in both tests. Memory was evaluated via the Hopkins Verbal Learning Test-Revised, with higher MOCA scores indicating greater levels of memory (range 0–12). The speed of processing was examined using the Trail Making Test A, with lower scores indicating greater levels of processing speed. Visuo-spatial ability was examined using the Brief Visuospatial Memory Test-Revised, with higher scores indicating greater levels of visuo-spatial ability (range 0–12). Language function was examined using a verbal fluency test for animal naming, where higher scores indicate greater levels of language. Executive function was assessed via subtests, including a 100-stimulus version of the Stroop color and word test, a digit span test, a space span test, and a picture completion test; higher scores indicate greater levels of executive function. The complete cognitive assessment took about 120 minutes and took place at DCRC-PKUIMH.

### Image pre-processing

The pre-processing of resting state fMRI data was carried out using Statistical Parametric Mapping (SPM12)^21^ and Data Processing Assistant for the R-fMRI (DPARSF) toolkit^22^. The main steps included: (1) dropping off the first ten EPI volumes; (2) temporal correction for slice acquisition; (3) spatial normalization into the MNI space based on transformation parameters derived from aligning T1 images to the MNI standard template using diffeomorphic anatomical registration through the exponentiated lie algebra (DARTEL) method; (4) resampling to 3-mm isotropic voxels and spatially smoothing with a 4 mm full width at half maximum Gaussian kernel; (5) regressing out the following nuisances from each voxel’s time series, including 24 head motion parameters, global signal, cerebrospinal fluid, and white matter time series and linear trend; (6) filtering the residual time series within a frequency range of 0.01–0.1 Hz for reducing the effect of low-frequency drifts and high-frequency noise.

### Functional brain network construction

Based on the pre-processed fMRI images, the overall functional brain connectivity construction process is shown in Fig.7. First, we used a sphere-based functional brain atlas (Power et al.^12^, 2011) to define 264 brain regions of interest (ROI) belonging to 14 communities (neural systems) in total. Then, the mean signals (time-series) were computed within spheres of fixed radius *r* (*r* = 5) around a sequence of voxels in *T* functional brain scans (*T* = 230). Finally, we computed the Pearson’s correlation coefficient across all pairs of time series to construct the brain connectivity matrix C and obtained the corresponding 3D visualizations for functional brain connectomes. For every patient in the pre-processed fMRI dataset, we computed the brain connectivity matrices for fMRI data obtained at baseline (week 0) and week 12. The averaged brain connectivity results before and after interventions are shown in Fig. 8, and the corresponding brain graphs exemplified for one patient are plotted in Fig. 9.

**Fig. 6.**
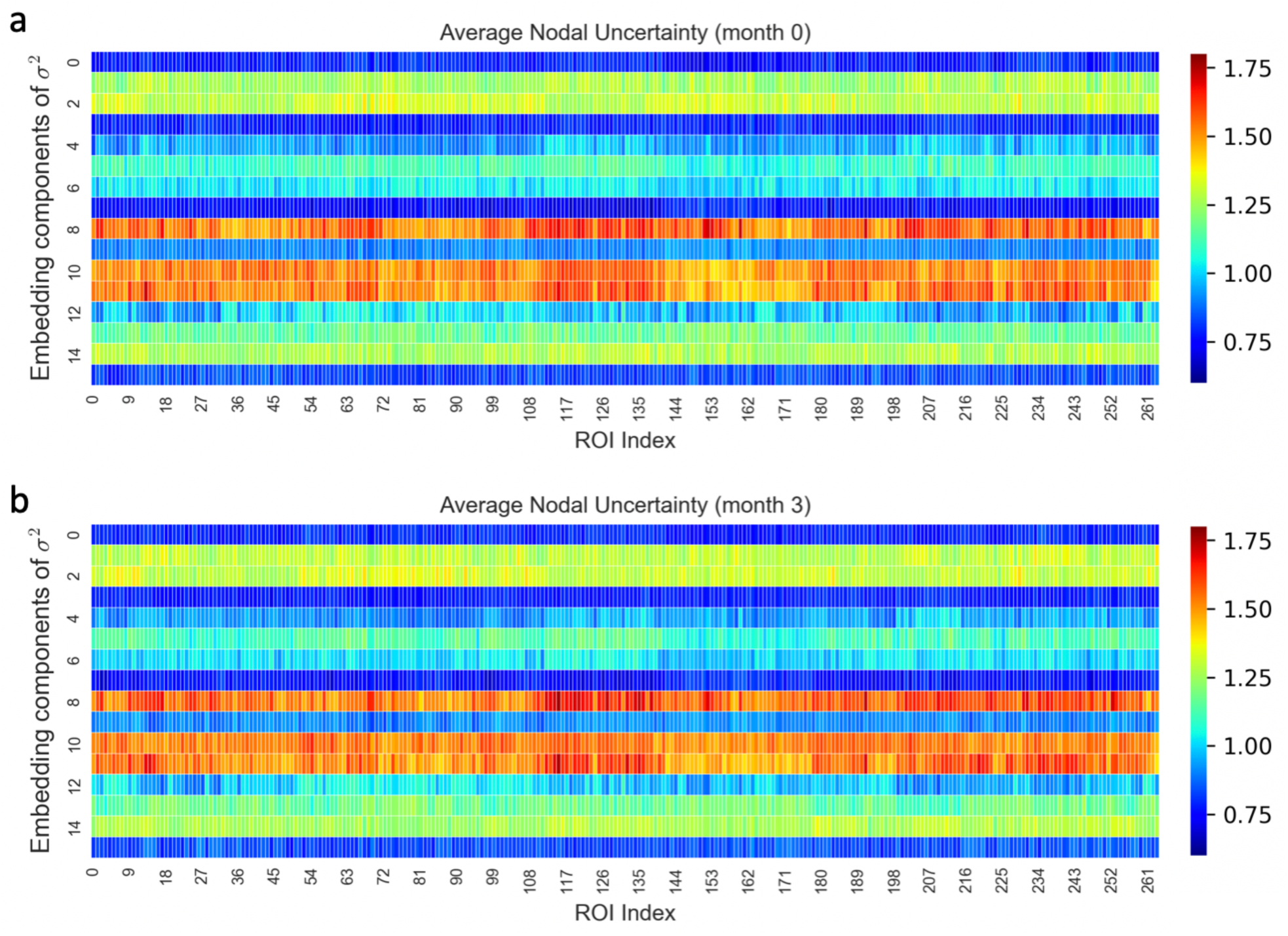
Uncertainty quantification using the MG2G approach. Average nodal uncertainty (variance-σ^2^) results for 12 patients before (a) and after intervention (b); (embedding size *L*=16).

**Fig. 7.**
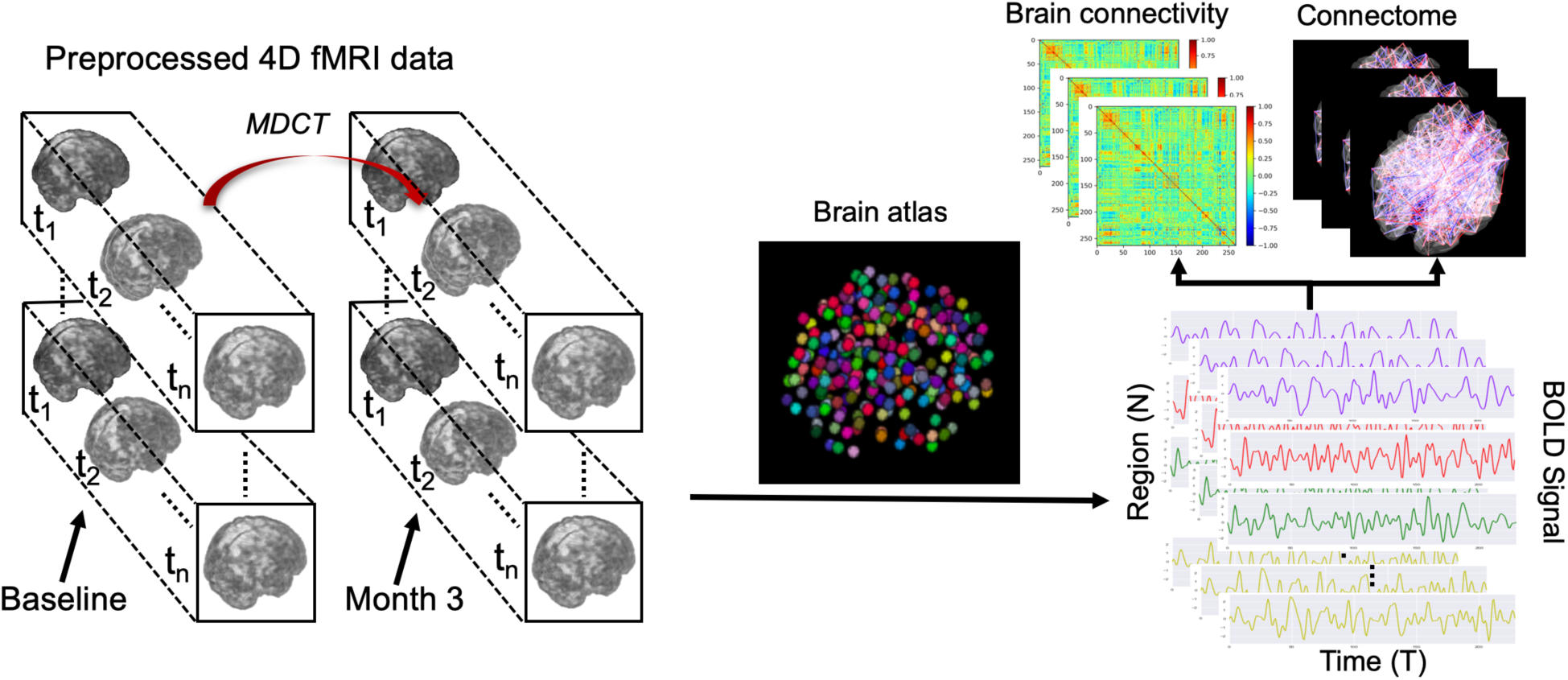
Illustration of brain connectivity construction workflow based on the Power et al. functional atlas^12^.

**Fig. 8.**
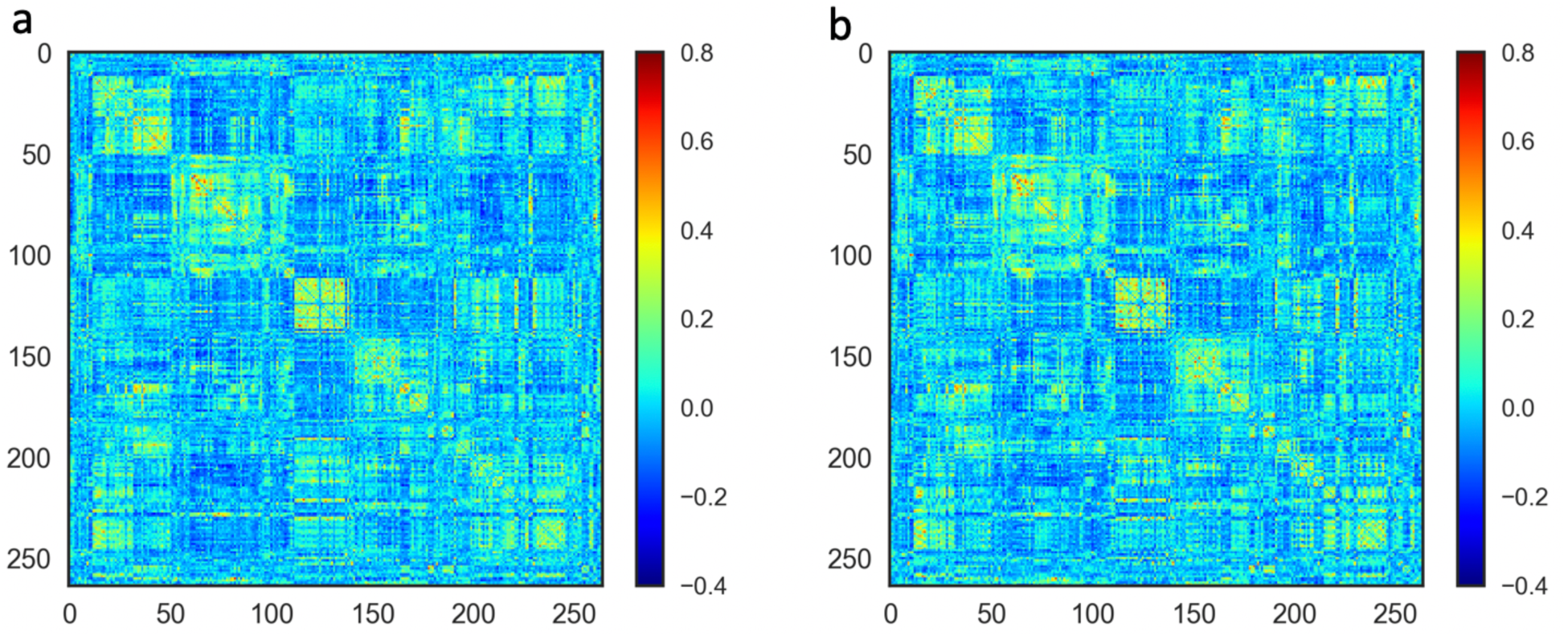
Brain connectivity matrices averaged across all 12 patients at baseline (a) and after 12-week MDCT intervention (b).

**Fig. 9.**
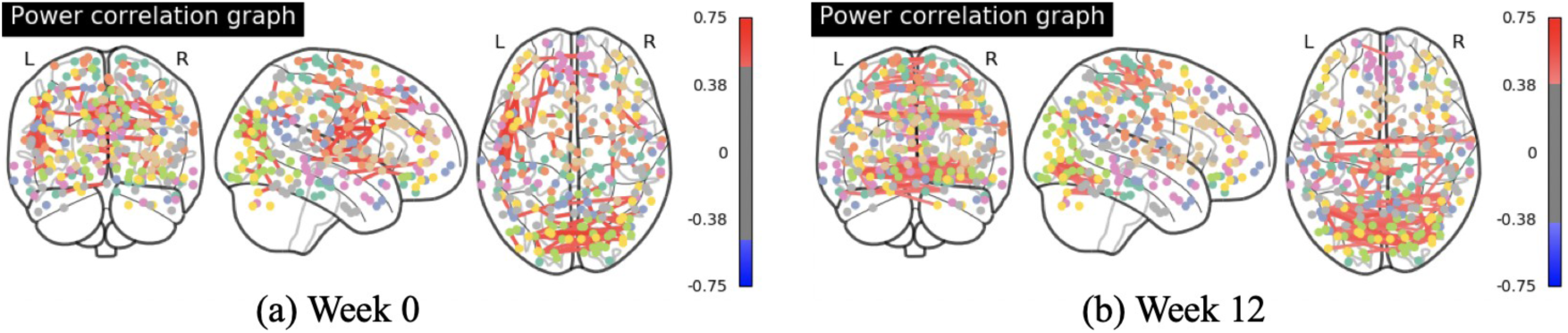
Graphical representations of brain connectivity for subject 1 at baseline (a), and after 12-week MDCT intervention (b).

### Multi-Graph2Gauss embedding approach for functional brain network analysis

Since functional brain networks calculated from our raw fMRI data were undirected and weighted, in our work we extended the Graph2Gauss method^15^ to a multi-graph Gaussian embedding (MG2G) prediction model for functional brain networks. This method can realize an unsupervised Gaussian projection learning from an original brain network into latent low-dimensional Gaussian distributions (Fig. 10).

**Fig. 10.**
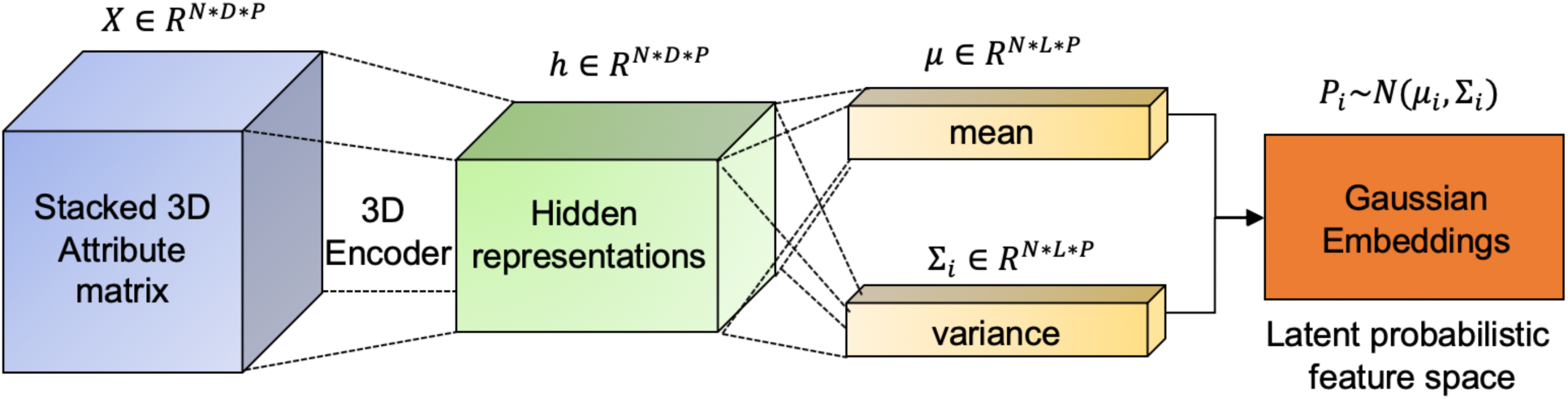
Main architecture of the proposed *MG2G* model for multiple human brain networks.

First, for each of the *P*=24 original brain connectivity matrices C, we obtained a thresholded adjacency matrix A by setting to 0 connectivity values below an empirical threshold (*t*=0.1). These thresholded matrices were then used to compute k-hop neighbors (*N*_*ik*_, *k* ≥ 2) for each brain region (node *i*) based on Supplementary Eq. 2, and sample valid triplet sets based on the obtained hops (Supplementary Eq. 3).

In addition, for each node *i*, we assigned as node attributes the *i*^*th*^ row vector of A. In other words, each node had attributes the connectivity profile (connection weights) across all the N nodes in the network. Consequently, brain graphs had N-dimensional node attributes, with the aim of subsequently compressing them to *L* dimensions via graph embedding.

In order to encode multigraph data jointly into the same space, we stacked all brain network’s edge attribute matrices into a 3D tensor. This yielded a 3D matrix *X* ∈ *R*^*N*×*D*×*P*^, where *N* denotes the number of brain regions in brain network, *D* denotes the dimension of attributes and is equal to *N* in our work, and *P* is the number of brain networks (Fig. 10). Matrix *X* as the input of the deep encoder architecture. We adopted a 3D encoder to learn a mediate latent representation, and in turn used it to output the means and variances of the final embedding Gaussian distributions for all patients. Furthermore, the optimization of deep neural networks was carried out by minimizing the square-exponential loss as shown in Supplementary Eq. 6.

In comparison with the basic principles of *Graph2Gauss* summarized in SI, our MG2G model for functional brain networks made three contributions: i) we made use a weighted (as opposed to binary) *symmetric adjacency matrix* to compute k-hop neighbors and triplet sets; ii) we added the connection weights as edge attributes to provide extra information for graph embedding, and iii) we extended the method to multigraph data.

As a metric of comparison and to capture the subtle differences before and after MDCT intervention, we made use of the 2-Wasserstein distance (Eq. 1) for quantitative evaluation of ROI-specific changes between encoded probabilistic Gaussian distributions with respect to each patients’ brain networks before and after interventions.

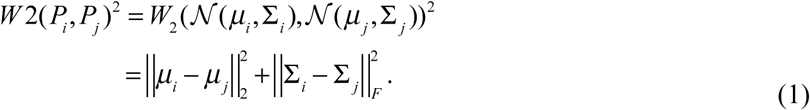

## Acknowledgements

We thank Prof. George Em Karniadakis of Brown University for fruitful discussions regarding the uncertainty quantification. MJX received partial support from NIH grant U01 HL1163232; ZJW, HFZ and HLW were supported by Beijing Municipal Science & Technology Commission (No. Z161100000516001, D171100008217007). MJX and DP were supported by a J-Clinic for Machine Learning in Health award at MIT.

## Author Contributions

MJX, DP and QZL conceptualized the research. ZJW, HFZ and HLW designed the MDCT intervention; HFZ collected the data; ZJW pre-processed the human imaging data; MJX developed and validated the method, ran the simulations and analyzed the data; MJX and DP wrote the manuscript; QZL and DP supervised the work; ZJW, HFZ and HLW revised the manuscript.

