## Supplementary Information for "Gaussian embedding-based functional brain connectomic analysis for amnestic mild cognitive impairment patients with cognitive training"

### SUPPLEMENTARY METHODS

#### Overview of Graph embedding methods

Graph embedding methods have gained a lot of attention in recent years since they can effectively project large-scale networks to a low-dimensional latent space, while preserving the intrinsic network topological properties. The obtained graph embedding can be used for downstream graph processing tasks - such as link prediction, node classification, and community detection - much more effectively, easily and with high computational efficiency compared to other more classical methods.

Recent survey studies ([1,2]) divided graph embedding techniques into three main categories: (1) matrix factorization-based approaches, (2) random-walk based approaches, and (3) deep learning-based approaches. For graph embedding methods, the primary challenge is to preserve the first-order and high-order proximity during graph embedding implementations. In order to tackle this problem, matrix factorization-based methods (e.g., GraRep [3] and HOPE [4]) construct a high-order proximity matrix based on transition probabilities and factorize it to obtain the node embeddings, but they are not easy to scale up for large network embeddings. Random walk-based methods (e.g., node2vec [5] and DeepWalk [6]) utilize different node neighbor set searching strategies through modified random walk paths to capture global and local proximities during the low dimensional point-vector embedding procedure. However, the major drawback of the aforementioned embedding approaches is the lack of capturing important "uncertainty" information for each node in the complex networks. Nodes with low degree contain fewer network connections, hence they have larger uncertainty than other nodes (or "entities" in the knowledge graph). Similarly, edges (relations) that link to more entities have a relative larger uncertainty than others. For the purpose of estimating *uncertainty information* for each node in the network, Gaussian embedding is uniquely positioned to derive and characterize embeddings in terms of means and variances in a space of multivariate Gaussian distributions. For example, Vilnis et al. [7] first proposed the word2Gauss method, which applied Gaussian embedding to map word types into a space of Gaussian distributions in order to model the uncertainty, entailment, and inclusion information of different word types in latent space. He et al. [8] proposed KG2G learning Gaussian embedding for latent knowledge graph representation. Moreover, Zhu et al. [9] proposed to apply deep variational network models in conjunction with the 2-Wasserstein distance and build a hybrid loss function to obtain Gaussian embeddings that preserve the transitivity in embedding space. Bojchevski et al. [10] employed a deep neural network model to learn node embedding as Gaussian distributions much more efficiently and robustly in latent embedding space for attributed and directed graphs. In particular, they developed an efficient way of predicting the effective dimensionality of the low-dimensional space (latent dimension) by monitoring during training the “uncertain" directions as the dimensions with the largest uncertainty, which are unstable and do not contribute to the low-dimensional embedded graph.

### Graph2Gauss embedding method

Compared with traditional graph embedding methods that project nodes into low-dimensional point vectors, deep neural network-based Gaussian embedding models, such as the *Graph2Gauss* model [10], can offer a very promising and novel approach for learning graph node representations (or “encodings") as a latent space of Gaussian distributions in an inductive and unsupervised manner. In the latent graph embedding space, each node is encoded as Gaussian distributions with two different learned vectors (mean and variance). The mean vector reflects the position of the node while the variance, usually constructed in two different shapes (diagonal or spherical), provides important uncertainty information. Specifically, the learned uncertainty provides information on two critical aspects: 1) correlation with neighborhood diversity*, i.e.* larger variance reveals more diversity in the node’s k-hop neighborhood, 2) ability to discover the intrinsic latent dimensionality of the complex graph, which is close to the number of ground-truth communities in the graph.

The graph embedding problem can be interpreted mathematically as follows. Given a directed/undirected graph $G = (A, X)$ with $V$ and $E$ the corresponding vertex and edge sets, $A$ denotes the (symmetric or asymmetric) adjacency matrix of size $N \times N$, and $X$ is the attribute matrix of size $N \times D$, the aim of Graph2Gauss is to project every node from a high-dimensional space into a latent space of multivariate Gaussian distributions. For instance, the embedding of node $i$ ($P_i$) can be represented by a $L$-dimensional Gaussian distribution with a mean vector ($\mu_i$) and a co-variance matrix ($\Sigma_i$) in diagonal shape, where $L \ll D$.

$$P_i \sim \mathcal{N}(\mu_i, \Sigma_i) \quad \mu_i \in \mathbb{R}^L, \Sigma_i \in \mathbb{R}^{L*L} \tag{1}$$

In order to obtain the latent Gaussian representation for every node in a graph, the main architecture of Graph2Gauss contains four main elements: 1) Unsupervised node representation learning based on a deep encoder; 2) Node embedding modeling as Gaussian distributions; 3) Energy (distance) estimation for pairs of nodes in the embedding space; 4) Gaussian embedding learning by minimizing the energy-based loss, i.e. employing the square-exponential loss for the optimization of hyper-parameters of the deep encoder. The specific workflow is illustrated in Supplementary Figure 1.

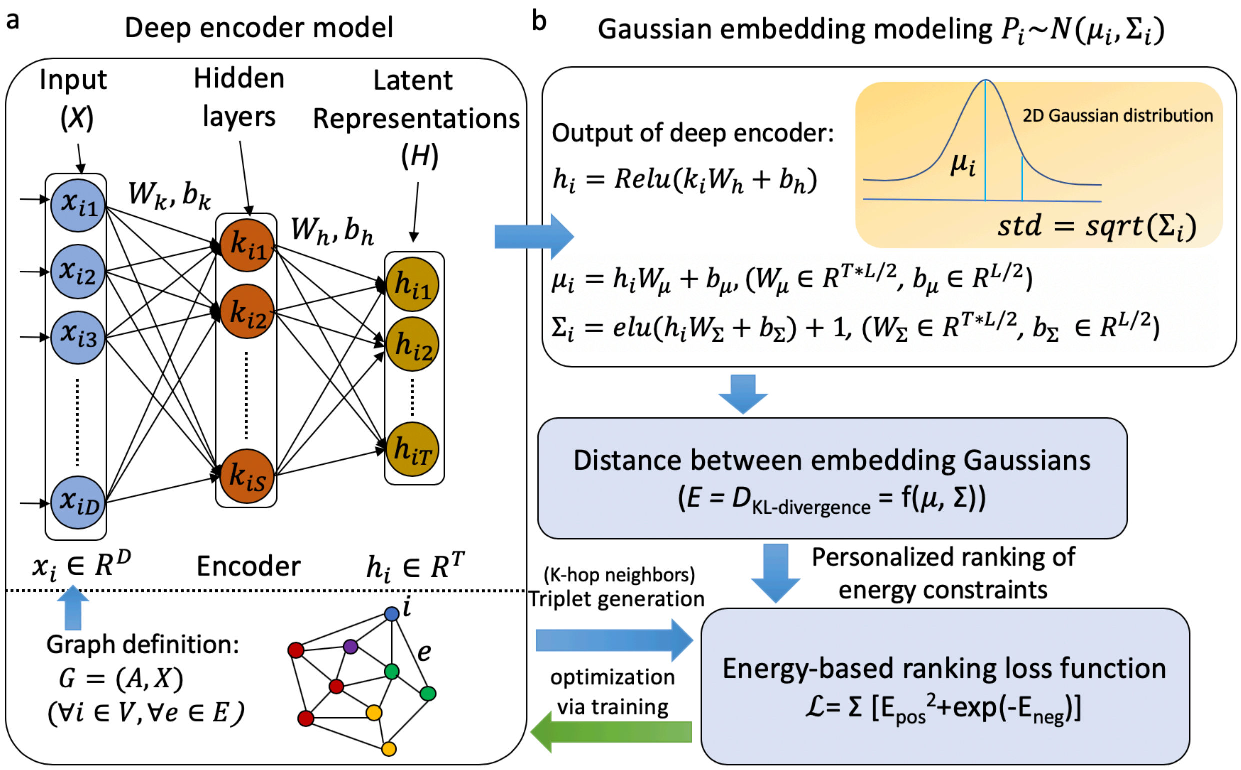


**Supplementary Figure 1. Illustration of the main workflow for the Graph2Gauss embedding approach**. a) The original graph and node attributes are input to a deep encoder, yielding the latent representations. b) The output of the deep encoder is fed to two neural networks for the mean and variance estimation. Optimization is based on an energy ranking loss function, which utilizes the k-hop neighborhood information.

Below, we provide details for some of the key components of the Gaussian embedding learning process.

*1) Node triplet generation*: Given a node $i$, its $k$-hop neighbors can be represented as $N_{ik}$ in Eq. 2, where $sp(i, j)$ denotes the shortest path between node $i$ and node $j$ (if $i$ and $j$ are not reachable, it returns ∞); $K$ is the maximum considered distance, usually $K > 2$ enables capturing high-order proximity. A triplet sample usually consists of anchor, positive, and negative nodes [11]; a set of valid triplets can be represented as in Eq. 3, with $j_k \in N_{ik}$, $j_l \in N_{il}$ and $k < l$. Thus, for the triplet $(i, j_k, j_l)$, node $i$ is more similar to node $j_k$ than node $j_l$, and the node pair $(i, j_k)$ denotes one positive pair, while the node pair $(i, j_l)$ denotes one negative pair, which are generated for the energy-based ranking loss construction. Moreover, a "node-anchored sampling" strategy [10] provides an effective way for triplet generation that can help reduce the computational complexity in a large graph.

$$N_{ik} = \{j \in V \mid i \neq j, min(sp(i,j), K) = k\} \quad (2)$$

$$D_t = \{(i, j_k, j_l) \mid sp(i, j_k) < sp(i, j_l)\} \tag{3}$$

<u>2) Network structure preservation via personalized ranking</u>: In order to capture the network structure properties at a multiscale level for graph embedding, personalized ranking of energy (similarity) constraints (Eq. 4) are imposed to the latent node embeddings, *i.e.* the respective energy (or distance) between embeddings of node $i$ and each node in its $k$-hop neighbors ("positive energy") is lower than the one between embeddings of node $i$ and its ($k$+1)-hop neighbors ("negative energy"). Here, $E(P_i, P_j)$ denotes the energy function between learned Gaussian distributions $(P_i, P_j)$ for nodes $i$ and $j$. The specifications of energy function can be seen below:

$$E(P_i, P_{k_1}) < E(P_i, P_{k_2}) < \ldots < E(P_i, P_{k_K}) \quad \forall k_1 \in N_{i1}, \forall k_2 \in N_{i2}, \ldots, \forall k_K \in N_{iK} \tag{4}$$

<u>3) Similarity quantification for embedding Gaussians</u>: The energy function ($E$) in Eq. 4 is used to measure the distance between two nodes' embedding Gaussian distributions $(P_i, P_j)$ and score the triplets. Here $(i, j)$ can be either positive node pairs or negative node pairs. Currently, most commonly used similarity metrics include the symmetric expected likelihood ($EL$), the Jensen-Shannon divergence ($JS$), the asymmetric KL-divergence ($KL$), and the $p$-th Wasserstein distance ($W_p$). Different from the first two metrics, $KL$ and $W_p$ can also handle directed graph embeddings while at the same time preserving the transitivity of nodes. The asymmetric "KL-divergence" energy is shown in Eq. 5, where tr(.) denotes the trace of a matrix, and det(.) denotes the determinant. Smaller $E$ represents that the nodes' embedding Gaussians are more similar or closer to each other.

$$\begin{aligned} E(P_i, P_j) &= D_{KL}(P_j \mid P_i) = \int_{x \in \mathbb{R}^n} \mathcal{N}(x; \mu_i, \Sigma_i) \log \frac{\mathcal{N}(x; \mu_j, \Sigma_j)}{\mathcal{N}(x; \mu_i, \Sigma_i)} dx \\ &= \frac{1}{2}[tr(\Sigma_i^{-1}\Sigma_j) + (\mu_i - \mu_j)^T \Sigma_i^{-1} (\mu_i - \mu_j) - L - \log \frac{det(\Sigma_j)}{det(\Sigma_i)}] \end{aligned} \tag{5}$$

<u>4) Energy-based ranking loss function</u>: Based on the aforementioned constraints on the respective energy between latent embeddings of adjacent $k$-hop neighbors of each node (*e.g.* 2-hop vs. 3-hop of node $i$), in order to learn a graph embedding that satisfies the constraints, we adopt an energy-based learning approach [12]. That is, we build an energy-based ranking loss ($\mathcal{L}$) over the sampled triplets ($D_t$) for penalizing the ranking errors, such that positive energy ($E_{ij_k}$) terms are always lower than negative energy ($E_{ij_l}$). $\mathcal{L}$ usually consists of two parts: positive node pair energy ($E_{ij_k}$) and negative node pair energy ($E_{ij_l}$). "Margin-based ranking loss" is a frequently used loss function for graph embedding learning, however, the margin has to be manually selected before training. Thus, the "square-exponential loss" [12] representing negative pair energy as an exponential term has better performance in penalizing the ranking error automatically; the specific formula is given in Eq. 6.

$$\mathcal{L} = \sum_{(i,j_k,j_l)\in D_t} (E_{ij_k}^2 + exp^{-E_{ij_l}}), k < l \tag{6}$$

### Example of uncertainty quantification

To illustrate how Graph2Gauss, and specifically our proposed MG2G method, estimates uncertainty, we plot Gaussian distributions for several nodes (ROIs) of one patient in Supplementary Fig. 2. The difference of two Gaussians is also a Gaussian with mean the difference of the two means and variance the sum of the two variances.

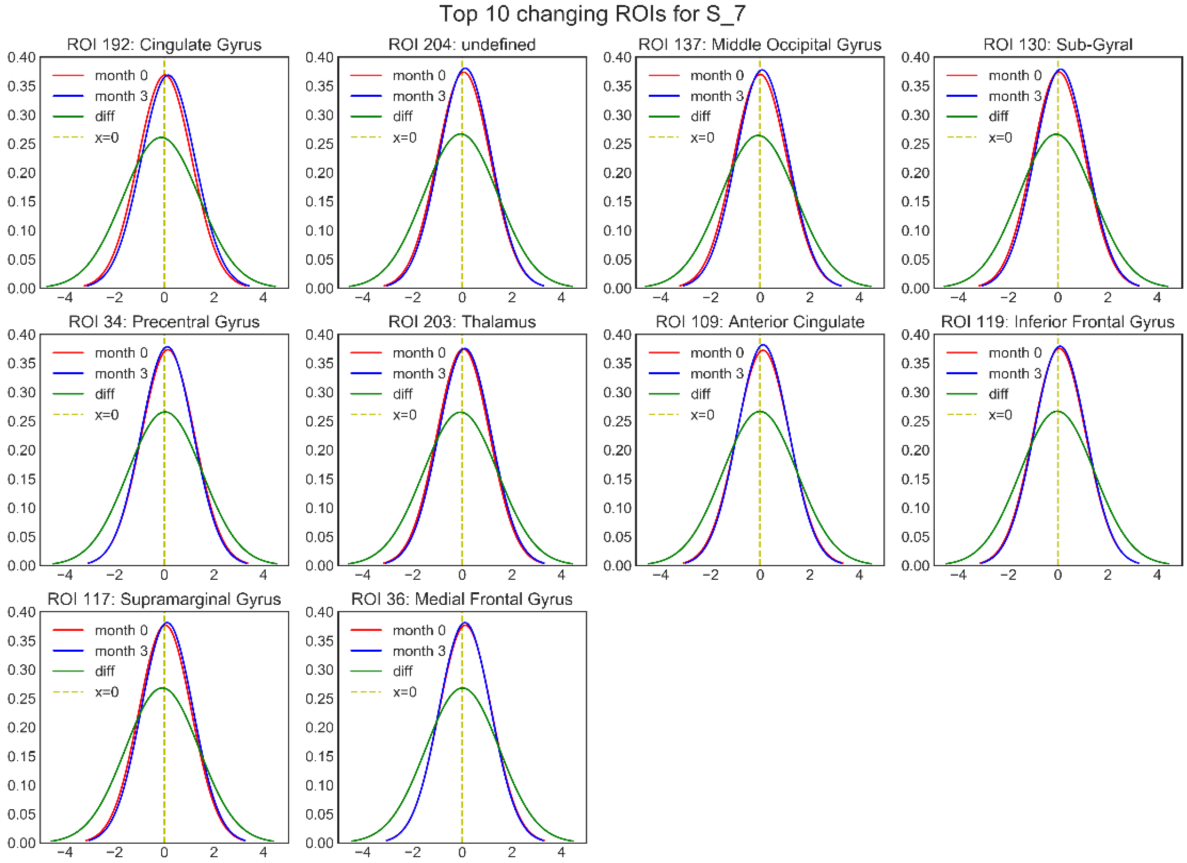


**Supplementary Figure 2. Gaussian distributions obtained from MG2G for one patient.** ROI-specific embedding Gaussian distributions before (red curve), after (blue curve) MDCT intervention, and the distribution of the difference (green curve). We only include the top-10 ROIs with the highest W2-distance.

### Comparison of MG2G versus node2vec

We evaluated the performance of our proposed MG2G method against node2vec in Supplementary Fig. 3, which shows system-level changes for each patient using MG2G, and Supplementary Fig. 4, which shows the corresponding results using node2vec. The output of node2vec is deterministic and hence we can only compare mean values with MG2G.

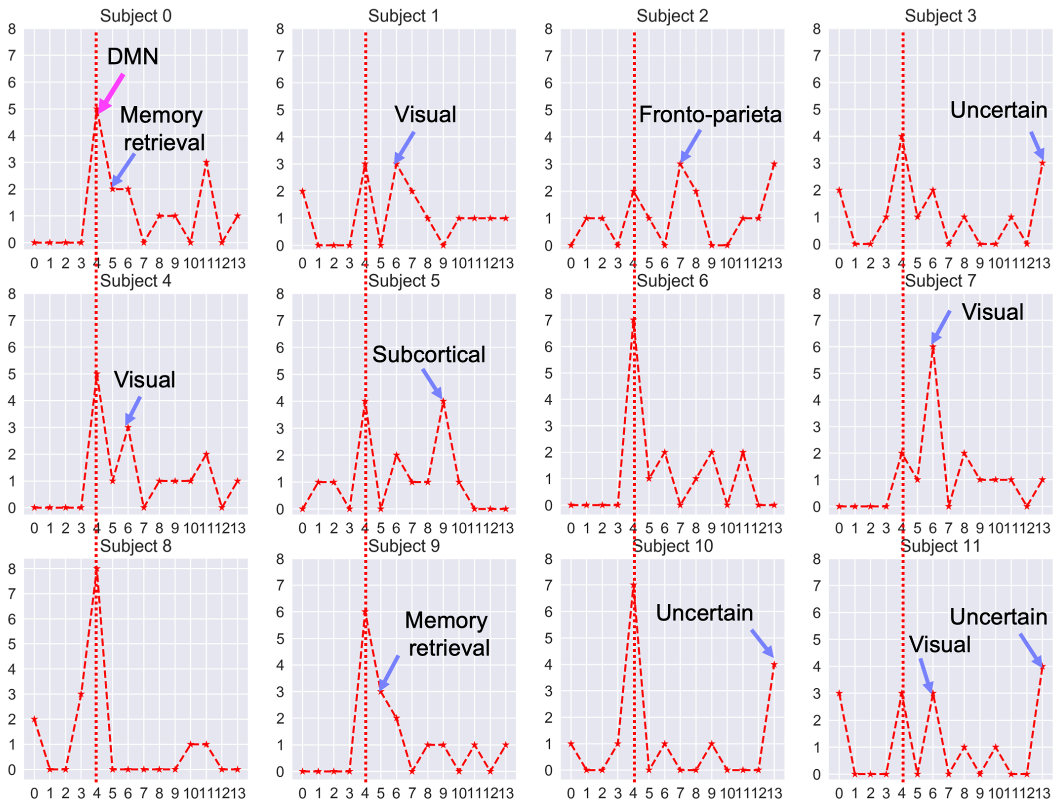


**Supplementary Figure 3. Quantification of system-level changes for all 12 patients before and after MDCT intervention based on MG2G**. The plots show the number of top-15 ROIs with the highest network alterations contained within different functional brain systems. System names same as in main text Fig. 3.

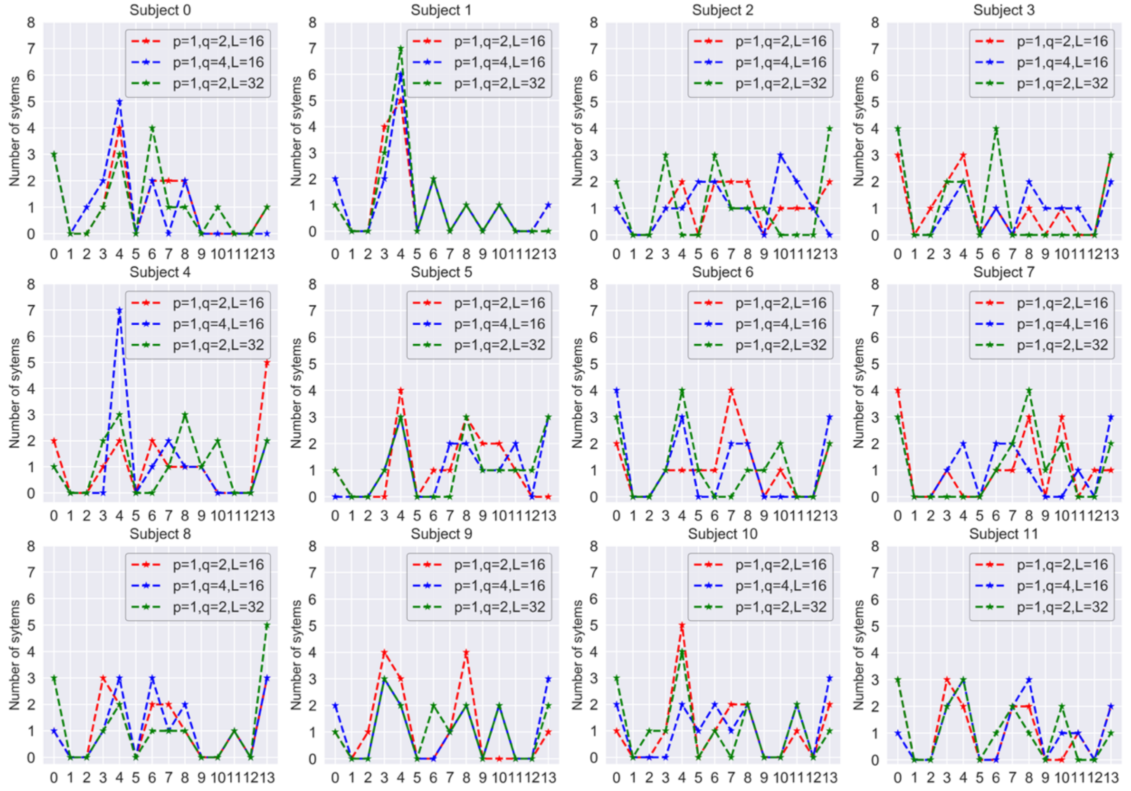


**Supplementary Figure 4. Quantification of system-level changes for all 12 patients before and after MDCT intervention based on different parameters of node2vec.** We tested performance with different embedding sizes (L = 16, 32) and p, q values in node2vec; these values control the neighborhood exploration in node2vec.

### Effect of k-hop neighborhoods

We evaluated the performance of MG2G for link prediction based on different k-hop neighborhoods (k=2, 3) in Supplementary Fig. 5. Results show that k = 2 is sufficient and there is no need to consider higher-order proximity measures.

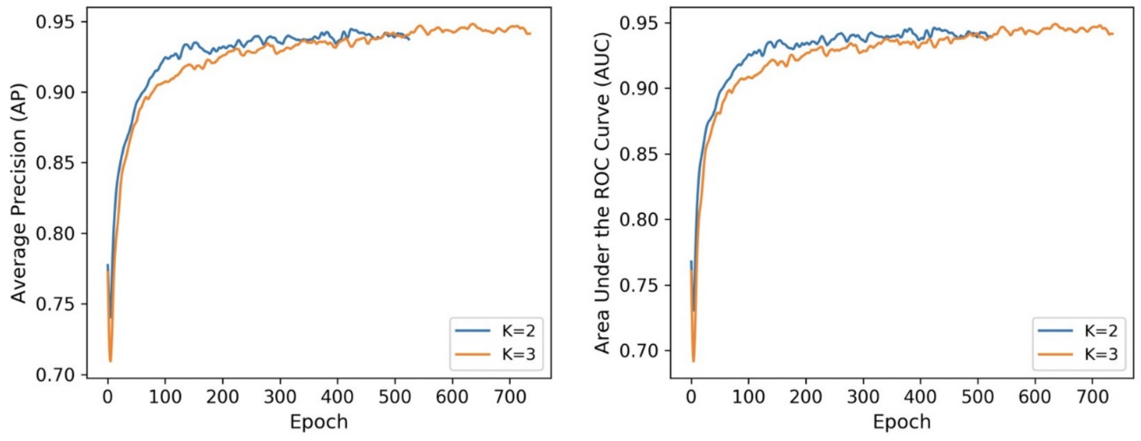

(a) (b)

**Supplementary Figure 5. Evaluation of link prediction performance using MG2G based on different k-hop neighborhoods.** (a) Average Precision vs. number of epochs, (b) AUC vs. number of epochs. In both cases, the embedding size (L) was equal to 16.

### Example of reorganization index (RI) estimation

We estimated the RI for the different ROIs comprising the neural system 'sensory/somatomotor hand' (SSH). Both ROIs (8 and 28) that achieved the highest RI values are located in the precentral gyrus of the right hemisphere. The majority of the ROIs had positive RI values, indicating extensive reorganization of the SSH neural system.

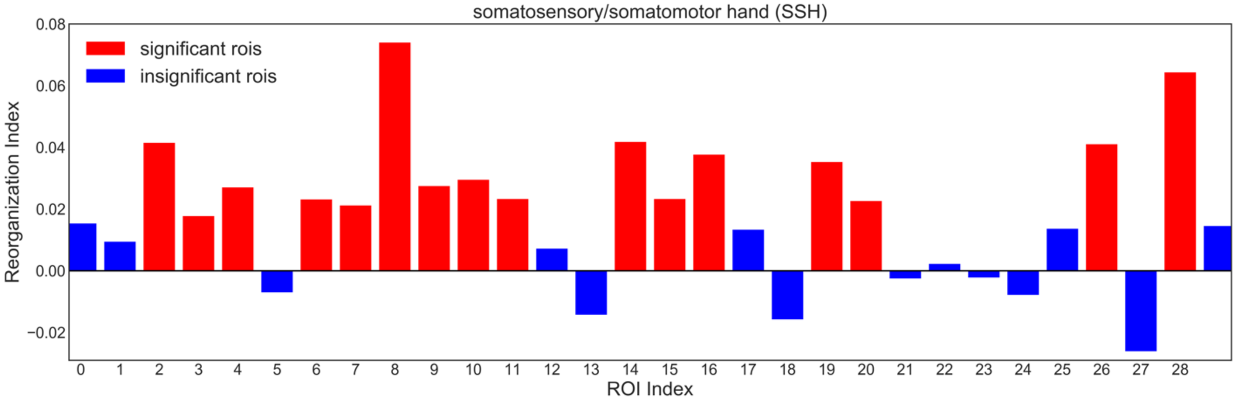


**Supplementary Figure 6. Reorganization index results for 30 ROIs in the '*somatosensory/somatomotor hand'* functional brain system**. A large number of ROIs had significant RI (red bars; $p<0.05$, FDR corrected). System name abbreviations same as in Supplementary Table 1.

### Brain Atlas Description

| NO. | ROI range | System | Abbreviation |
|---|---|---|---|
| 1 | 0-29 | Sensory/somatomotor Hand | SSH |
| 2 | 30-34 | Sensory/somatomotor Mouth | SSM |
| 3 | 35-48 | Cingulo-opercular Task Control | CoTC |
| 4 | 49-61 | Auditory | Audit |
| 5 | 62-119 | Default mode | DMN |
| 6 | 120-124 | Memory retrieval | MemRt |
| 7 | 125-155 | Visual | Vis |
| 8 | 156-180 | Fronto-parietal Task Control | FpTC |
| 9 | 181-198 | Salience | Sal |
| 10 | 199-211 | Subcortical | SubCt |

| | | | |
|---|---|---|---|
| 11 | 212-220 | Ventral attention | VenAtt |
| 12 | 221-231 | Dorsal attention | DorsAtt |
| 13 | 232-235 | Cerebellar | Cerebl |
| 14 | 236-263 | Uncertain | Uncert |

**Supplementary Table 1. Functional brain atlas information.**
